# Detection of Amyotrophic Lateral Sclerosis (ALS) via Acoustic Analysis

**DOI:** 10.1101/383414

**Authors:** Raquel Norel, Mary Pietrowicz, Carla Agurto, Shay Rishoni, Guillermo Cecchi

**Author notes:** Deceased 24 May 2018 (Shay Rishoni).

## Abstract

ALS is a fatal neurodegenerative disease with no cure. Experts typically measure disease progression via the ALSFRS-R score, which includes measurements of various abilities known to decline. We propose instead the use of speech analysis as a proxy for ALS progression. This technique enables 1) frequent non-invasive, inexpensive, longitudinal analysis, 2) analysis of data recorded in the wild, and 3) creation of an extensive ALS databank for future analysis. Patients and trained medical professionals need not be co-located, enabling more frequent monitoring of more patients from the convenience of their own homes. The goals of this study are the identification of acoustic speech features in naturalistic contexts which characterize disease progression and development of machine models which can recognize the presence and severity of the disease. We evaluated subjects from the Prize4Life Israel dataset, using a variety of frequency, spectral, and voice quality features. The dataset was generated using the ALS Mobile Analyzer, a cell-phone app that collects data regarding disease progress using a self-reported ALSFRS-R questionnaire and several active tasks that measure speech and motor skills. Classification via leave-five-subjects-out cross-validation resulted in an accuracy rate of 79% (61% chance) for males and 83% (52% chance) for females.

## 1. Introduction

ALS is a progressive, incurable, neurodegenerative disease that leads to decreased muscle function resulting in problems with movement, breathing, swallowing and speech. The disease is referred to as limb onset or spinal ALS when the first symptoms appear in the arms and legs, and bulbar ALS when it presents with speech or swallowing difficulty. Patients with limb onset ALS maintain speech much longer than those with bulbar onset; however, 80% of all ALS patients experience dysarthria, or unclear, difficult speech articulation [1]. On average, speech remains adequate for about 18 months after the first bulbar symptom appears [2]. Experts measure disease progression with the ALSFRS-R score, which measures various abilities known to decline as the disease progresses. We envision instead the use of automatic speech analysis as a proxy for disease detection and progression. In this paper, we evaluate speech features which characterize speech deterioration in ALS using the Prize4Life Israel dataset, and we validate them in the context of machine models which recognize disease presence.

The characteristics of dysarthria in ALS include hypernasality, hoarseness, strain, slow rate, imprecise articulation, monotone quality, reduced volume, longer stop closures, longer vowel duration, and smaller vowel space [3]. In addition, acoustic analysis has shown that ALS speech exhibits longer stop closures, longer vowel duration, deviant F0, jitter, shimmer, and unstable voice quality/phonation [1]. Nasality fluctuation also occurs in dysarthria, when velar control is affected [4]. The specific symptoms of dysarthria will vary from patient to patient, depending on the structures affected [1]. Research suggests that ALS dysarthria is distinct from other causes, such as frontotemporal dementia [5]. Outside the realm of ALS, dysarthria is also multifaceted, affecting intelligibility and articulation, with varying severity. Prior work on automatic detection of generic dysarthria has explored these different facets and severity levels using a variety of analytic techniques [6]–[11].

Progression rate of dysarthria also varies, and the change in speaking rate and intelligibility over time can identify slow and fast disease progressors [12]. Speech impairment may even begin close to 3 years prior to ALS diagnosis [13], offering the potential for monitoring high-risk subjects (e.g., familial ALS and head concussions) early. Clinical assessment often requires the reading of specially-designed passages designed to elicit dysarthria or apraxia of speech [14], and our work uses such passages. Several clinical assessment scales exist for dysarthria, but comparative assessment tools for the scales themselves do not exist. The dataset used in this study includes ALSFRS -R[2] scores for each recording. These ALSFRS-R speech scores, however, do not reflect the specific attributes of dysarthria, are subject to bias via self-report, and have poor granularity to characterize the many variations of ALS speech [1].

Prior work using speech in the detection of ALS typically uses kinematic sensors to model articulation [15]-[16], analyzes acoustic patterns in speech, or measures prosodic elements such as vowel duration or speaking rate. Wang *et al*. [17] investigated lip and tongue articulation data with acoustic data simultaneously, and found that acoustic data alone could function as well as acoustic data with lip and tongue data in an SVM regression of acoustic features using a balanced set of speakers with respect to intelligible speaking rate. The features were selected from the OpenSMILE [18] set, and the same features were applied to both males and females. A second study explored classification of the ALS condition using the same features applied to SVM and DNN classifier [19]. Alternate approaches used formant trajectories to classify the ALS condition [20], correlated formants with articulatory patterns [16], or used fractal features [21]. Our work extends prior efforts in that it 1) functions on non-ideal, non-studio data collection conditions, as are likely to occur in at-home or in-clinic recording scenarios, 2) considers an order of magnitude larger set of speakers and samples than the prior studies, 3) considers the different acoustic characteristics of males and females, and selects features which optimize analytics by gender, 4) requires fewer features than prior work to perform at least as well, in part because models are gender-specific (a single feature suffices for male speakers), 5) uses a baseline set of sentences and paragraphs recommended by speech therapists for evaluation of dysarthria in ALS, and 6) functions with training data collected across an unbalanced range of widely-varying stages of disease progression as represented by widely-used ALSFRS-R scores, as is likely to occur with data collection in the wild. Accuracy of our SVM classifiers exceeds comparable prior results, reported in literature.

Our approach is synergistically positioned with the coming mobile devices and systems which support longitudinal data collection and analysis of ALS and other conditions. One such system is the ModelTalker voice banking pipeline [22], and another is the Prize4Life ALS Mobile Analyzer [23], which monitors patient disease progression from home. The eventual goal is deployment of a reliable system which uses unbiased monitoring of speech quality as a proxy for condition progress monitoring, to enable prompt action in response to actual or predicted changes in patient condition. Our contributions toward this goal include 1) a gender-optimized, selection of features which characterize ALS dysarthria, 2) improved accuracy detection rates in SVM classifiers, 3) a reduced feature set size for simplicity and computational efficiency, and 4) a data analysis process and machine model which is tolerant to recordings not made in studio conditions.

## 2. Database curation and analysis

The data originated from the ALS Mobile Analyzer [23], which was designed, developed, and deployed by the non-profit organization Prize4Life Israel [24]. Its purpose was to enable the digital monitoring of disease progression in patients, without requiring them to leave home. The continued monitoring and tracking of patients will result in a growing database of objective measurements which may, in turn, help researchers find biomarkers of disease progression, facilitate the search for effective disease treatments, and ultimately, accelerate progress toward a cure. We focus on the resulting speech recordings here, which contain scripted speech collected from ALS patients and their caregivers (controls). The subjects had to read 3 sentences or 1 paragraph in English at their own pace at a preferred location. Note that not all the subjects were native English speakers, and not all subjects read all the suggested sentences.

The dataset required curation because many of the original recordings were taken in uncontrolled conditions and were unsuitable for analysis. Files which contained less than 2 seconds of speech, contained less than 2 syllables, (measured using nuclei syllables detection [25]), or contained loud background noise were removed from consideration. Next, the surviving files were filtered to removed silence and remaining sections of non-speech noise. The resulting corpus contained approximately 100 utterances each from male and female speakers, nearly evenly divided across controls and patients. The number of speakers in each condition, however, were unbalanced (27 female patients, 40 male patients, 30 female controls, 26 male controls), and we compensated for this in the analysis. Given that mean age for controls was about 9 years younger than patients, we regressed out age using the data of the controls and applying the correction to all. Regressing age-corrected features to age confirmed that the age effect had been eliminated.

The resulting dataset had one notable bias. The female portion of the dataset had a distribution of patients across the spectrum of severity, as measured by ALSFRS-R scores; but the male portion of the dataset was biased toward healthier patients whose conditions had not progressed as far as many of the female patients’ conditions had progressed. This means that the average male voice sample was closer to normal speech, and that the male model had fewer representative examples to characterize the more severely affected voices. Given this bias, and physiological differences, we chose to estimate independent models for males and females.

**Fig. 1:**
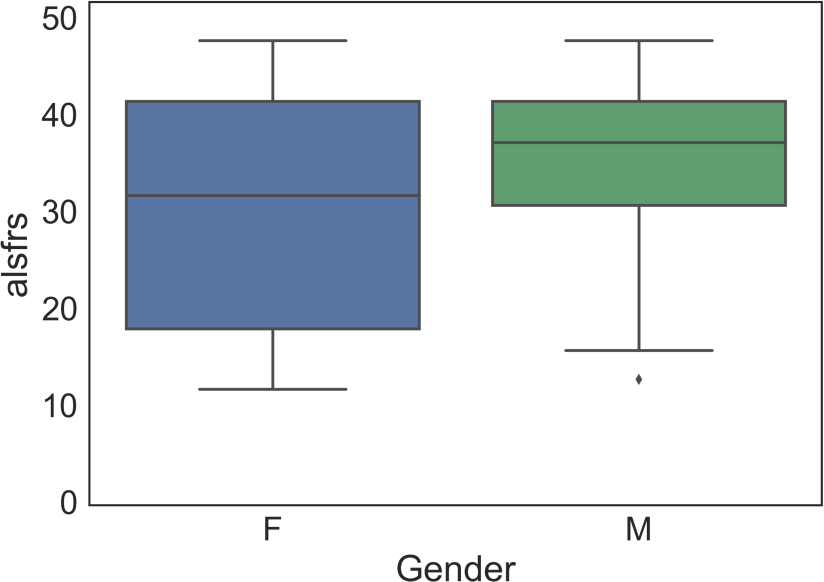
*Total ALSFRS-R score distribution for Female/Male (F/M) subjects in the analysis. Most of the male patients’ ALSFRS-R scores ranged between 33-42, while most of the female patient’s scores ranged between 18-42. A lower ALSFRS-R score represents a more advanced disease progression*.

## 3. Methods and statistical analysis

The goal is the correct classification of sound files from ALS patients and controls, for both males and females. To characterize the speech of the participants in the study we first extract features using openSMILE [18], perform feature selection using a t-test, build models using standard classifiers, and cross validate leaving 5 subjects out.

### 3.1 Feature extraction

The openSMILE [18] toolkit provides a baseline set of pitch, energy, waveform, auditory, spectral, voice quality, and other important features for consideration. It includes many popular analytic techniques, including Cepstral analysis variants, formants, and RASTA-PLP, and provides support for both low-level descriptors (LLDs) and summary analytics (thousands of potential features). We adopted the ComParE13 [26] subset (60 msec frames and a 10-second hop) for baseline analysis, given its successful use in the Paralingual Challenge. Tables 1 and 2 show the most significant features for females and males, respectively, and Figure 3 characterizes the most informative single features providing the greatest separation between conditions.

**Table 1:** *Top significant features for female speakers*

| Feature Name (Female) | p-val |
| --- | --- |
| mfcc_sma[3]_linregc2 | 1.91E-11* |
| mfcc_sma[1]_linregc1 | 6.20E-10* |
| mfcc_sma[3]_quartile3 | 1.02E-09* |
| audSpec_Rfilt_sma[25]_percentile1.0 | 4.70E-09* |
| mfcc_sma[3]_qregc3 | 7.82E-09* |

**Table 2:**
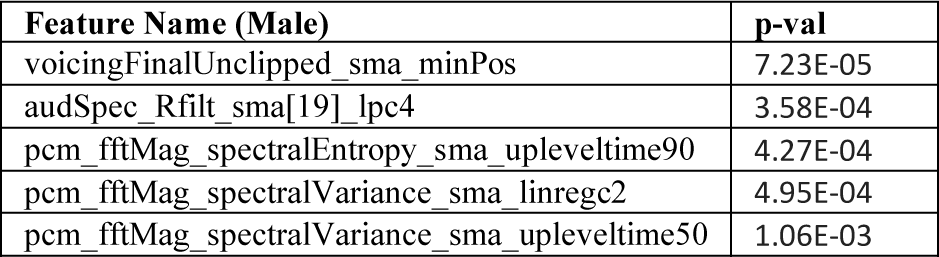
*Top significant features for male speakers*

**Fig 3:**
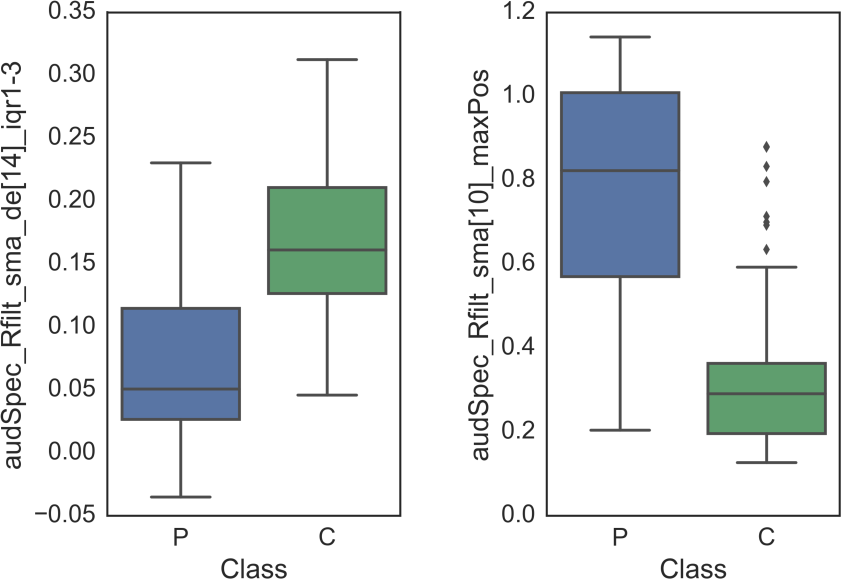
*Distribution of values for most informative feature after age correction for female (left) and male subjects to discriminate patients vs. controls*.

### 3.2 Statistical analysis and feature selection

The ComParE13 openSMILE feature set contained thousands of features, so we check whether the features for inclusion in our models are statistically significant. Since multiple comparisons were performed, we applied false discovery rate (FDR) correction at q<0.05. Tables 1 and 2 below show the top statistically-significant candidates for the female and male models respectively. Note that all of the female features passed the FDR correction; however, none of the features from the male model survived the FDR correction criteria for statistical significance. Note that we used gender-specific features and created gender-specific models, since male and female vocal tracts are acoustically different.

Note that the selected male and female features are quite different. The only similarity in the set of top significant features are the “audSpecRfilt_sma” features, where the coefficient differences correspond to characteristic differences in the male and female vocal tracts. Beyond this similarity, MFCC coefficients characterize female voices, and spectral changes, the male voices.

### 3.3 Classification and validation

After feature standardization (μ=0, σ=1) we performed a two-nested leave-subject-out cross validation approach using linear support vector machines (SVM) classifier for performance estimation and parameter selection, respectively. Feature selection was performed via univariate selection (two-sample t-test) in the internal cross-validation procedure. The resulting features were used to train nine off-the-shelf classifiers (Decision Tree, Linear Support Vector Machine, Linear Discriminant Analysis, Logistic Regression, Logistic Regression with L1 norm, Naive_Bayes, Nearest Neighbors, Random Forest, Support Vector Machine with elastic net regularization). Classifier performance was validated via a leave-five-subject-out cross-validation approach, with classifier parameters being selected via an internal 5-fold cross-validation. In other words, we used a nested cross-validation scheme to calculate classification error rates and optimal parameter values.

## 4. Results

Figure 2 below shows the results of the representative Linear SVM classifiers for males and females. The male classifier had a 79% accuracy rate (precision=0.78, recall=0.76) and the female classifier had 83% accuracy rate (precision=0.86, recall=0.78).

**Figure 2:**
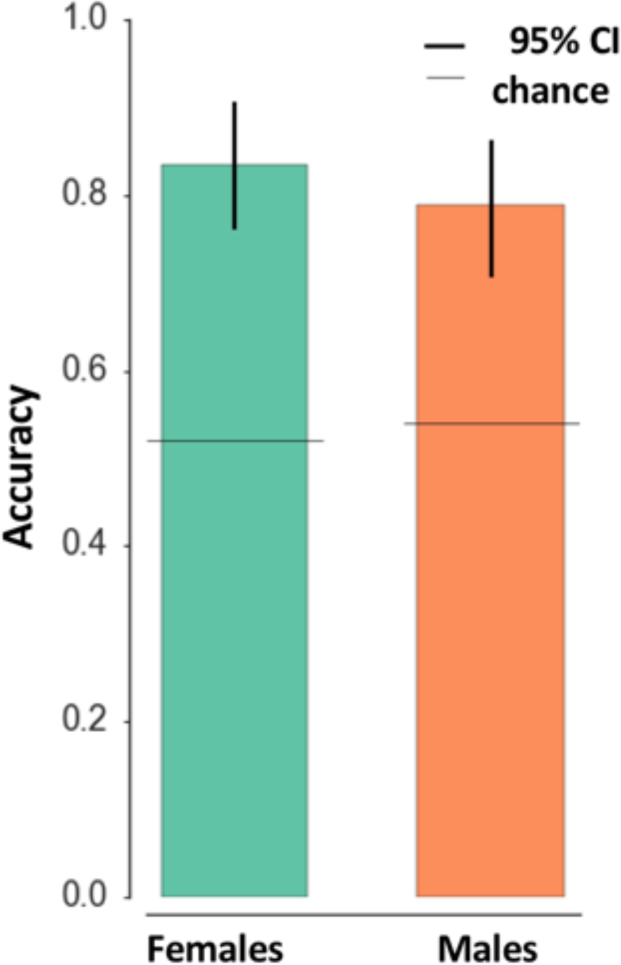
*Classification performance for Linear SVM classifiers, using leave-five-subject-out cross validation. Confidence intervals at 95% are marked with black vertical lines. Chance probability is calculated and displayed with a horizontal black line. Results obtained with leave-five-subject-out and ten-fold cross validation surpass chance probability*.

Not all features performed equally well. For males, a single feature could separate conditions with 79% accuracy. For females, the top 15 features could separate conditions with 78% accuracy. Figure 3 characterizes separation characteristics for the top male and female features. It is interesting to observe that the most informative feature for males is “audSpec_Rfilt_sma[10]_maxPos,” and the most informative corresponding feature for females is “audSpec_Rfilt_sma_de[14]_iqr1-3”. This implies RASTA filtering of the audio spectrum, smoothed, in different regions corresponding to gender. RASTA filtering highlights changes, or “edges,” in the sound as part of its process. For a visual analog, image a picture with lines drawn around each object and lines highlighting shadows, color changes, etc. This filtering not only highlights the sonic edges, but it also suppresses spectral components which vary at rates different from the typical rate of change in speech. This result suggests that spectral changes in ALS and non-ALS speech are characteristically different.

## 5. Discussion

The dataset used in this study presented challenges. The data did not originate from a clinical trial or contrived study, and therefore could not be expected to provide balanced data. This was seen in the difference in age between controls and patients, and in the unbalanced representation and distribution of patients across disease severity in males. We corrected the age bias via age correction across the features, but could not create data which did not exist to balance the representation of disease severity. The male model did not perform as well as the female model, probably because of this imbalance. In the future, larger sample sizes may help balance the training set with real data; otherwise artificial balancing techniques such as synthetic minority over-sampling technique (SMOTE) could be used.

The data was also not collected in laboratory conditions, but was instead recorded in the wild, often in noisy environments. The noisiest data, including recordings with music, loud crowd noise, other speakers, and loud noise events had to be discarded or cleaned. In the future we hope to perform the analysis with a larger cohort of subjects, and hope to provide concrete advice and procedures for clean data collection to avoid heavy background noise levels. In addition, we advise the recruiting of control subjects which match the age profile of the patients. The advantage of using this kind of dataset, however, is that the resulting models will function on data collected in the wild, which is the level of robustness required for deploying mobile symptom tracking tools.

The utterances in this dataset were very limited. Most of the utterances were one of three short sentences, or a longer paragraph. Although these passages were designed to elicit abnormal speech in patients who had dysarthria, they were not representative of the range of normal speech, and not all subjects uttered all sentences. This imbalance could also have introduced biases into the dataset which could have affected model performance. The literature reports that ALS patients have smaller vowel space than non-ALS patients [1], but we were unable to detect this in our dataset, possibly because the sentences were short and did not contain enough exemplary vowels. We suggest a wider range of scripted utterances, along with the elicitation of unscripted speech.

The subjects in our dataset came from many countries and spoke many different native languages; many had obvious accents. They were all asked to read the same English sentences or paragraphs. Language differences could have generated confounding artifacts such as prosodic variation carried over from the native tongue, differences in vowel pronunciations and durations, and slower speaking rate. Slower speaking rate, especially, is a known result of both ALS and of speaking an L2 language. We purposely did not analyze speaking rate because of this potential confusion and instead searched for markers which would be accent independent.

In spite of the dataset limitations, our linear SVM model performance exceeded comparable SVM models. We believe that system performance can be improved by using deep learning methods, particularly on larger datasets.

The patient assessment procedures could also be improved. The most commonly-used severity assessment scale, the ALSFRS-R, limited our analytic options, particularly with respect to exploring regression. The granularity of measurement of speech fluency was not sufficient for our analytic goals; and many subjects who exhibited obvious dysarthria symptoms rated a “4,” the highest rating on the scale. These subjects were indistinguishable on paper from the controls or patients who did not yet exhibit discernable speech difficulty. Furthermore, the frequency of measurement was not consistent across subjects and did not support longitudinal analysis. In addition, recording the onset site (bulbar or spinal) would be helpful to analysis given the implications on prognosis [27] and speech deterioration [28]. Finally, the patients were monitored by self-reporting answers from a questionnaire, and these self-reported numbers were used to calculate the ALSFRS-R score. Self-reporting is inherently biased.

## 6. Conclusions

We demonstrated successful recognition of ALS and non-ALS speech on a dataset collected in the wild with no special equipment. The resulting solution used off-the-shelf feature extraction (openSMILE) and classification methods (linear SVM) on scripted sentences designed for speech assessment (accuracy=83% for females and 79% for males). This end result was a gender-optimized solution with improved performance over comparable linear SVM classifiers. We found that we could use a small feature set size (even a single feature for males) for simplicity and computation efficiency. Finally, the result produced a data analysis process and machine model shown to be tolerant to in-the-wild recording conditions.

## 7. Acknowledgments

We are deeply thankful to the patients and caregivers that participated in using the ALS Mobile Analyzer App and provided data. We thank Neta Zach for multiple very helpful discussions.

